# *Infino*: a Bayesian hierarchical model improves estimates of immune infiltration into tumor microenvironment

**DOI:** 10.1101/221671

**Authors:** Maxim Zaslavsky, Jacqueline Buros Novik, Eliza Chang, Jeffrey Hammerbacher

**Affiliations:** Department of Genetics and Genomic Sciences, Icahn School of Medicine at Mount Sinai, New York, NY 10029; Department of Microbiology and Immunology, Medical University of South Carolina, Charleston, SC 29425

## Abstract

Robust quantification of immune cell infiltration into the tumor microenvironment may shed light on why only a small proportion of patients benefit from checkpoint therapy. The immune cells surrounding a tumor have been suggested to mediate an effective response to immunotherapy. However, traditional measurement of immune cell content around a tumor by immunohistochemistry, flow cytometry, or mass cytometry allows measurement of only up to a few dozen markers at a time, limiting the number of immune cell types identified. Immune cell type abundances may instead be estimated *in silico* by deconvolving gene expression mixtures from bulk RNA sequencing of tumor tissue. By measuring tens of thousands of transcripts at once, bulk RNA-seq provides a rich input to algorithms that quantify cell type abundances in the tumor microenvironment, affording the potential to quantify the states of a greater number of immune cell types (given adequate training data). Here, we first review existing methods for deconvolution and evaluate their performance on synthetic mixtures. Then we develop a Bayesian inference approach, named *infino*, that learns to distinguish immune cell expression phenotypes and deconvolve mixtures. In contrast to earlier approaches, *infino* accepts RNA sequencing data, models transcript expression variability, and exploits the relationships between cell types to improve deconvolution accuracy and allow interrogation from the level of broad categories to the level of finest granularity. The resulting probability distributions of immune infiltration could be applied to numerous questions concerning the diverse ecology of immune cell types, including assessment of the association of immune infiltration with response to immunotherapy, and study of the expression profile and presence of elusive T cell subcompartments, such as T cell exhaustion.

## Introduction

While the tumor microenvironment holds many secrets about the state of the tumor and whether the patient will respond to therapy, this region is difficult to interrogate. As part of the immune system’s response to cancer, immune cells of many different kinds infiltrate the area around a tumor. Numerous studies have demonstrated that the immune cells present in the tumor microenvironment are associated with patient prognosis -- an association stronger than even the prognostic value of the standard tumor TNM staging system, which rates cancers from stage I to stage IV [1]. Additionally, there is some evidence to suggest that the tumor microenvironment may modulate a patient’s response to checkpoint blockade [2–4]. Measuring the contents of the tumor microenvironment could help explain the differential response to checkpoint therapy, which has a response rate between approximately 15% and 30% [5–7]. Finally, the task of understanding the immune cell profiles that make up an environment as diverse as the tumor microenvironment can shed light on the vast ecology of immune cell types, including giving us new abilities to examine phenotypes of interest. For example, exhausted T cells are likely to be implicated in the response to checkpoint blockade [8]. But the presence of such a T cell compartment in the tumor microenvironment has until now been difficult to establish, undermining attempts to study the aggregate association of this cell type with clinical conditions.

The difficulty of interrogating the tumor microenvironment is attributable to a manual measurement protocol that is extremely low throughput and to computational alternatives that fail when exposed to complex mixtures, which are common in the region. The predominant methods to identify the immune cells within the tumor microenvironment involve significant manual intervention. These approaches begin with manual tissue section preparation. Fluorescence-activated cell sorting (FACS) may be used to separate immune cell types by their distinct surface markers. Alternatively, samples may be stained with antibodies to differentially color different cell types by immunohistochemistry (IHC). A pathologist must manually examine the images and count the cells of each variety [9].

Computational alternatives instead manipulate a proxy source of data to estimate infiltration. Recent “infiltrate quantification” methods exploit the fact that clinical tumor biopsies often undergo bulk RNA sequencing. Several computational approaches estimate the relative abundance of many immune cell types in a bulk gene expression mixture extracted from the tumor microenvironment. Indeed, they perform well for many mixtures (Figure 1a). These methods are unique because they are highly multiplex: they make use of tens of thousands of features. Detailed deconvolution of many cell types is difficult by multicolored IHC, which in common practice can stain with up to only seven dyes at a time [4,10]. While FACS and CyTOF allow more markers to be measured simultaneously [4,11], they remain dwarfed by RNA sequencing, which measures thousands of genes at once -- therefore capturing a fuller expression profiles within cell types, as well as affording the potential to examine many more cell types. Moreover, bulk RNA sequencing remains significantly cheaper and more common that single cell RNA sequencing. RNA-seq is commonly performed on tumor tissue for other purposes, so an immune infiltrate quantification method that functions on bulk RNA-seq data can easily be applied to samples for whom RNA-seq has previously been acquired for other analyses [12].

**Figure 1:**
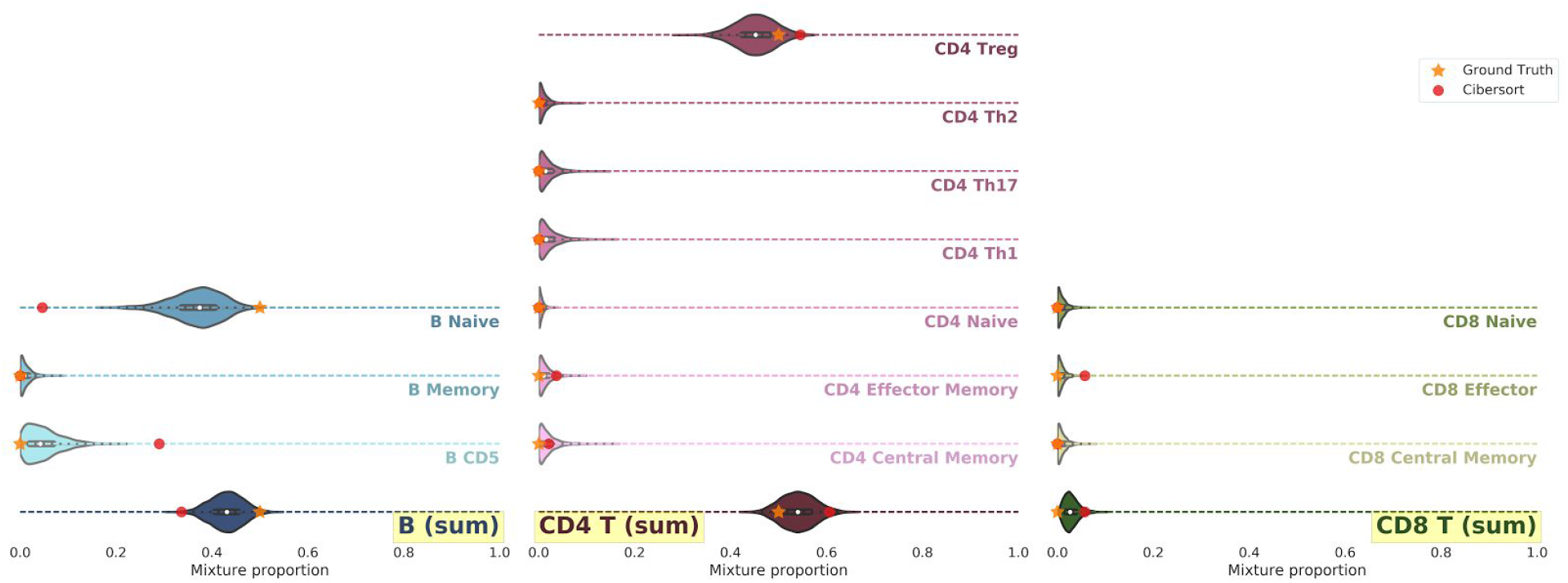
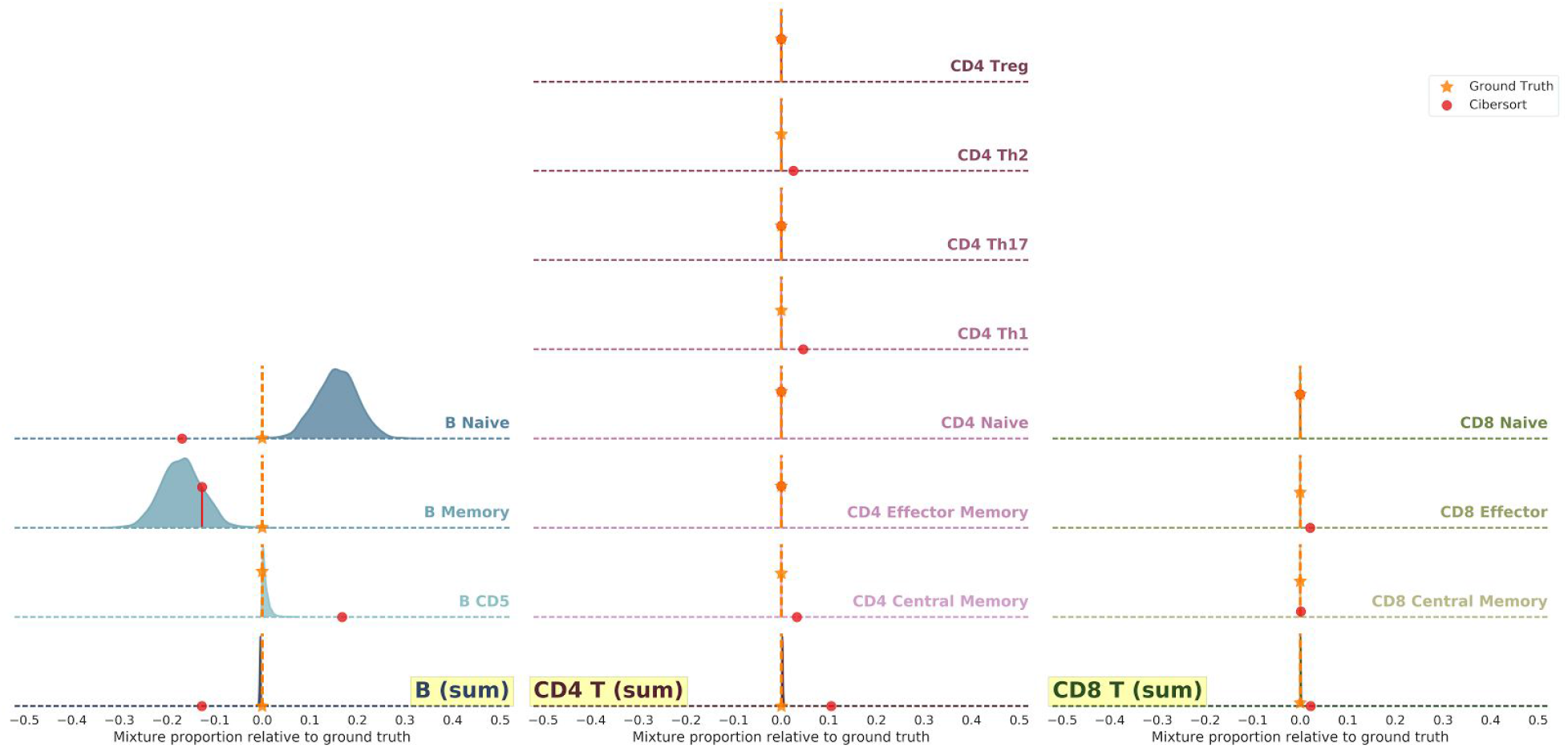
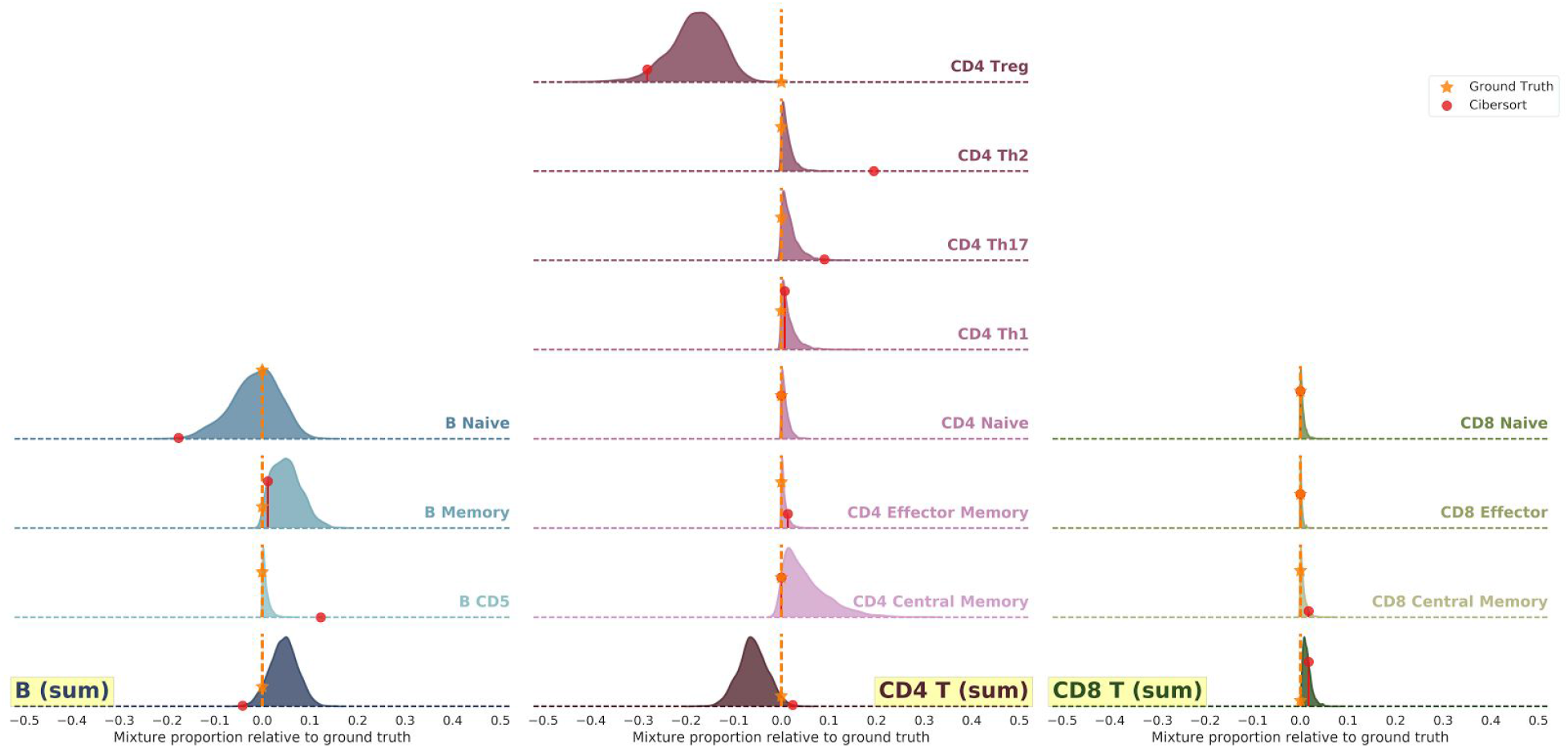
Deconvolution results by *infino* and *Cibersort* for several synthetic RNA-seq mixtures. The ground truth mixture weights are represented by yellow stars. *Infino*’s estimates of the fractional contribution of each subset to the mixture are shown as probability density distributions. Cibersort’s estimates are overlaid as red circles. Aggregated estimates at the B, CD4+ T, and CD8+ T cell supertype level are also illustrated in darker colors (highlighted rows). (a): Successful deconvolution by *infino* of a simple example, consisting of a 50%-50% mixture of a naive B cell sample and a CD4+ regulatory T cell sample. Cibersort’s diagnostics: p<0.01, RMSE=0.45. (b): Unsuccessful deconvolution of a more complex example, consisting of a 50%-50% mixture of a naive B cell sample and a memory B cell sample. Deviation of estimated mixture weights from ground truth is shown. However, *infino’s* aggregated scores demonstrate high confidence and precision (see highlighted rows). Cibersort’s diagnostics: p<0.01, RMSE=0.34. (c): Another complex example, a 25%-75% mixture of a naive B cell sample and a CD4+ regulatory T cell sample, reveals that *infino* underestimates regulatory T cell abundance. Deviation of estimated mixture weights from ground truth is shown. Cibersort’s diagnostics: p<0.01, RMSE=0.49.

To characterize the blended overall expression mixture collected from the tumor microenvironment -- which also includes stromal and tumor cells -- several groups identified marker genes whose differential expression is characteristic of certain immune cell types [13–19]. We seek to evaluate the marker gene approach to deconvolution. Since many immune cell types are remarkably similar in their gene expression profiles, we demonstrate that the strict criteria to identify marker genes can extract genes whose biological function does not appear unique to their associated cell types. For example, we performed gene ontology enrichment analysis on genes labeled as T cell markers only by the IRIS method [13], extracting biological pathways over-represented in the gene list relative to their expected background frequency [20,21]. The thirteen most significantly overrepresented gene ontology terms (evaluated at the significance threshold p < 0.001) were all related to mitotic nuclear cell division, a process not unique to the particular behavior of T cells. While [19] demonstrate how stricter criteria for marker genes identification can improve deconvolution, the authors note an important limitation: some immune cell types had no marker genes pass the threshold.

Other approaches first compute a representative expression profile for each immune cell type from purified cell populations, then model a test mixture as a linear combination of these reference profiles, solving for the mixture weights that produce the sampled mixture [22,23]. However, we identified shortcomings in the methodologies by which these methods extract representative profiles. First, we observed that the state-of-the-art method Cibersort [23] excludes relevant information, modeling only a point estimate of a transcript’s expression in each cell type, as opposed to its full distribution (Figure 2a). We then found that Cibersort fails to separate challenging mixtures of similar cell types, producing estimates with perfect confidence despite a high error rate (Figure 1b). Finally, many existing methods were designed only for data from microarrays, a technology that has been largely superseded by RNA sequencing in cancer research and clinical practice. As a result of these incomplete solutions to the problem of immune infiltrate quantification, no conclusive test of the predictive value of infiltration for response to checkpoint therapy has yet been performed, to our knowledge. In addition to shedding light on this important question, improving immune infiltrate deconvolution would enable identifying expression signatures for phenotypes of interest, such as exhausted T cells. As more granular immune cell subsets are identified -- particularly via single cell RNA sequencing -- and as expression profiles for these more detailed immune-cell subsets become available, it will be crucial for a deconvolution method to separate very similar cell types well and to facilitate a summary of inferences at any level of the cell-type hierarchy. In turn, this will require more interpretable and nuanced ways to evaluate the quality of a deconvolution.

**Figure 2:**
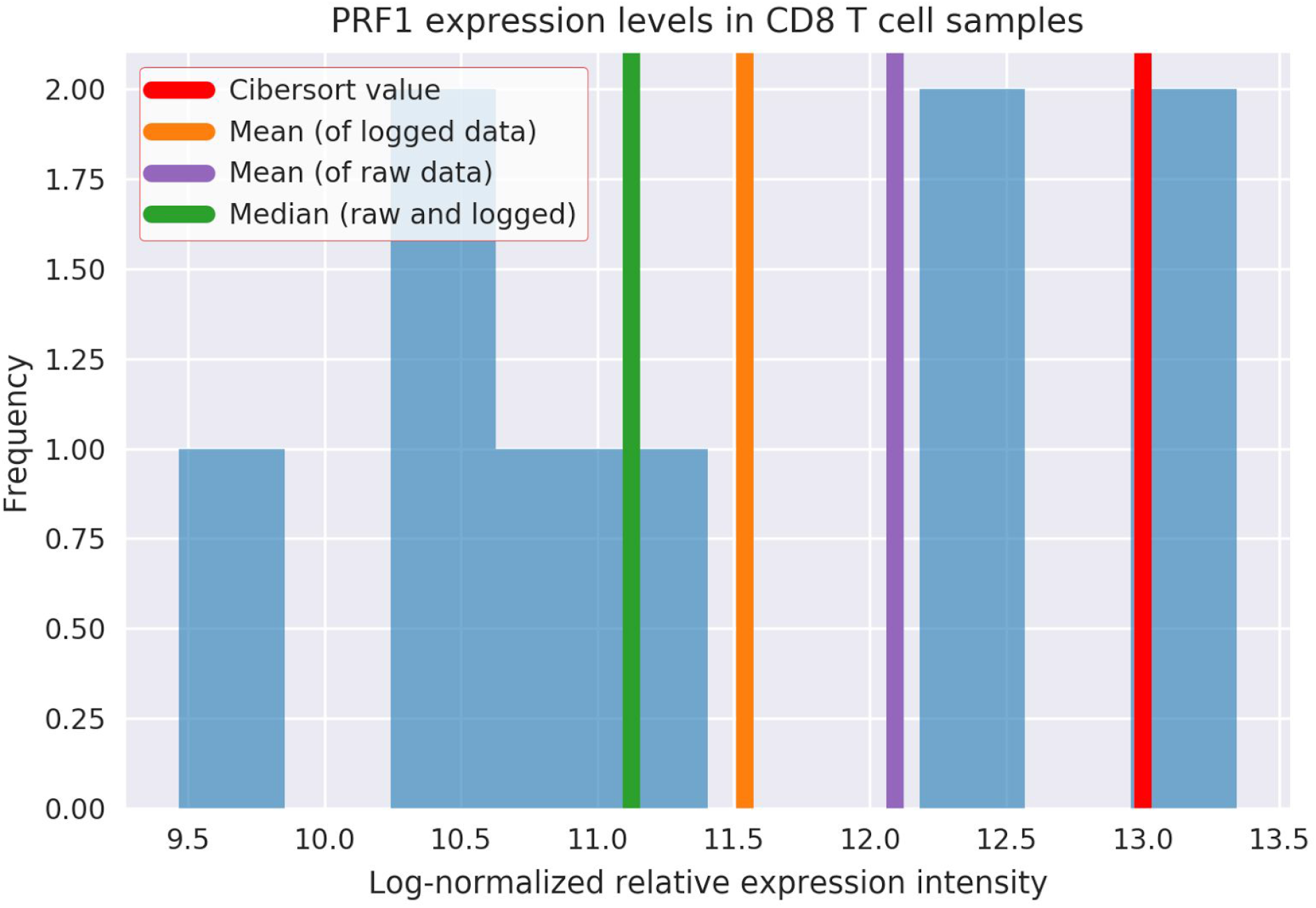
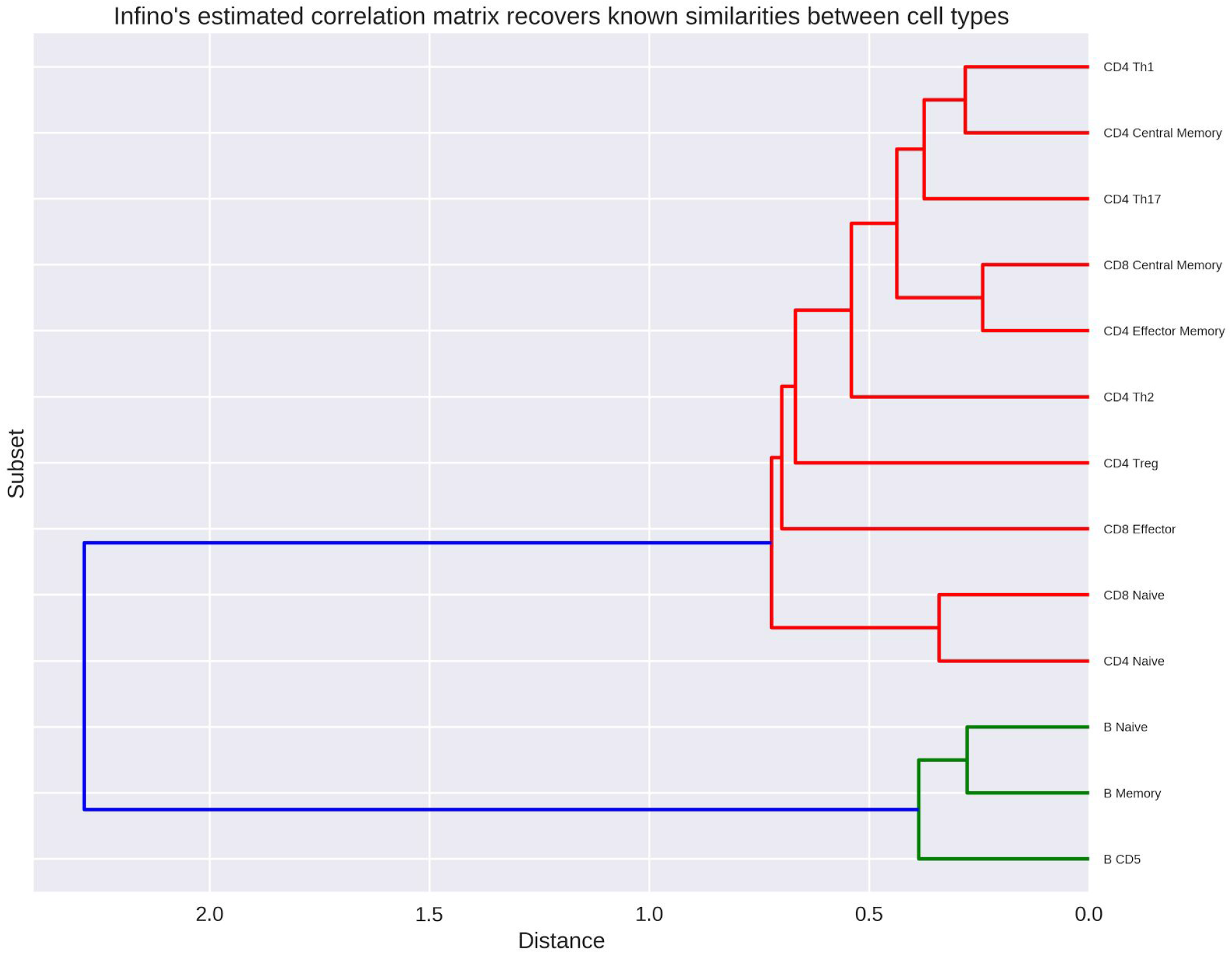
(a) Observed variance in expression levels of perforin 1 (pore forming protein), which has been suggested to be a CD8 T cell marker gene [14]. Histogram: nine microarray expression samples [GEO accession GSE22886, 13,GEO accession GSE6740, 24]. Cibersort’s representative expression level (from LM22) of this gene in CD8 T cells is overlaid in red. For comparison, summary statistics of interest, computed before and after log-normalization was applied to the microarray expression intensities, are also shown as vertical bars. The noticeable differences between the range of observed expression intensities, the summary statistics, and the value chosen by Cibersort suggest that a single point estimate may not be representative of transcript expression across samples of the same cell type. (b) Hierarchical clustering dendrogram created from the correlation matrix *infino* estimated from purified cell populations.

## Results

In this investigation, we refine the core statistical methodology for bulk tumor expression deconvolution to leverage additional information, including the natural hierarchy of immune cell subtypes. We introduce *infino* (“infer infiltrate expression phenotypes”), a new method that accepts clinical RNA sequencing data of gene expression in a patient sample and produces probability distributions of the abundance of many immune cell types. By applying Bayesian inference to this problem, we enable a clear representation of our model’s uncertainty in the deconvolution of a gene expression mixture from many cells into mixture weights of 13 cell types. We show below that while *infino* performs comparatively to previous infiltrate quantification methods on common mixtures, this Bayesian model enables analysis of complex cases for the first time, thanks to rich diagnostics in the form of probability distributions over its estimates to indicate the uncertainty in deconvolution.

Three key principles differentiate *infino* from existing infiltrate quantification methodologies. First, we model the process by which a mixture is generated from individual underlying cells with varying gene expression. In doing so, we encode the gene expression distributions of each individual cell type, rather than extracting point estimates to represent a certain cell type’s expression profile. This is in contrast to earlier approaches; for example, we observed that the state-of-the-art method Cibersort discards the variability between samples from similar and different contexts (Figure 2a). As a result, *infino* captures the variability in every immune cell type’s gene expression and returns posterior probability distributions as output, which provide clearer diagnostics than the point estimates produced by earlier methods.

Second, we exploit the relationships between cell types to improve our deconvolution results. Existing approaches attempt to deconvolve complex gene expression mixtures directly into the abundances of naive B cells, memory resting B cells, and so on. In contrast, we recognize that naive B cells and memory resting B cells, for instance, have extremely similar gene expression profiles. We perform deconvolution with knowledge of the relationships between cell types: naive B cells and memory resting B cells are both subtypes of a “B cell” supertype. Therefore, we model the shared characteristics of all B cells with high certainty, as B cells are much easier to distinguish from T cells than naive B cells are from memory resting B cells. Furthermore, we identify the deviations from the master B cell gene expression distributions that make a certain subtype unique. By encoding information about the “hierarchy” of immune cell types, we can evaluate the uncertainty in *infino’s* results at any deconvolution depth on-demand. For example, in the case of a particularly challenging mixture, as in Figure 1b, *infino* may report sufficient confidence in the abundances of B cells and T cells, but may note low confidence in its further deconvolution into B cell compartments and T cell compartments. Such granularity in the reporting of results is unprecedented for immune infiltrate quantification; to our knowledge, all existing approaches ignore these cell-type relationships, flatten the hierarchy, and return a deconvolution result at the level of the most granular and nearly indistinguishable cell types. While it is possible to aggregate these low-level cell type estimates to a higher level of the hierarchy, no existing method provides confidence metrics to accompany such a rollup. Moreover, *infino* learns these relationships directly from the data, as opposed to following a pre-configured arbitrary set of cell type relationships (Figure 2b).

Finally, *infino* accepts input data from RNA sequencing, which is more commonly performed today in the research setting than microarray measurement of gene expression. Meanwhile, several existing approaches only accept microarray input data.

When making predictions at the finest level of granularity, *infino* performs comparably to existing approaches. In testing on simple synthetic mixtures, all approaches estimate mixture weights with low error (Figure 1a). When tested on complex synthetic mixtures, though all approaches have high uncertainty or error in their estimation (Figure 1b), the advantages of applying Bayesian inference to immune infiltrate quantification become clear. In particular, *infino* reports its low confidence in the form of a wide confidence interval for the most granular level of cell type subsets. The robust diagnostic of evaluating the standard deviation of *infino*’s probabilistic estimates provides a clear understanding of deconvolution performance.

In this challenging mixture case, the advantages of incorporating information about the relationships between cell types also become clear. While all approaches struggle to form low-error, high-confidence estimates of mixture weights at the finest level of granularity, only *infino* produces robust estimates at higher levels of the hierarchy. When *infino*’s estimates are aggregated to the B cell, CD4 T cell, and CD8 T cell groups, we observed lower uncertainty and very accurate estimates for challenging mixtures (Figure 1b). Indeed, *infino* automatically recovers biological facts when the model learns relationships between cell types directly from the data, distinguishing clearly between the CD4 T cell, CD8 T cell, and B cell supertypes (Figure 2b). As a result, this modeling approach enables a user to interrogate complex mixtures at increasingly fine levels of granularity to an acceptable level of prediction uncertainty.

In our experimentation with synthetic mixtures, several cell types appeared to be particularly challenging to deconvolve. Very similar cell types, like naive and memory B cells, are notoriously difficult to separate (Figure 1b). These are indeed the cell types estimated to have the most similar expression patterns (Figure 2b). *Infino*’s aggregation capability can rescue the deconvolution of these mixtures and provide robust estimates at a higher level of the hierarchy. But the model also underestimates the abundance of regulatory T cells (Figure 1c), a pattern that deserves further exploration.

## Discussion

Understanding the differential response to cancer immunotherapy motivated our study of immune infiltrate quantification. We can apply our new approach to clinical data to test the association between immune infiltration and response to immunotherapy. Because our approach is entirely computational and requires no manual scoring, it can produce a sample size large enough to accurately test the prognostic significance of infiltration and address this key question in cancer immunotherapy. In particular, we plan to apply *infino* to data gathered from multiple clinical trials of checkpoint blockade therapy, producing infiltrate predictions for each patient’s tumor microenvironment from bulk tumor expression data alone. First, we will assess the association between infiltration and immune activation -- a sanity check, as we expect a strong correlation. Second, we would examine whether infiltration levels separate nonresponders from responders, which would be an intriguing and meaningful result. Then we could also examine other clinical variables, like survival, to refine our understanding of the ramifications of immune infiltration.

These conclusions suggest several future directions for refining the *infino* approach. First, there are opportunities to adjust the features of the Bayesian mixture model that we apply to the task of mixture deconvolution. Incorporating tissue-specific priors could prevent our underestimation of regulatory T cell content. We could include other sample-level covariates, such as adjustments for batch effects related to particular data sources or for the tissues of origin, which can lead to the observed variability in expression profiles. Additionally, we can consider modeling cell surface markers, which may be shared between different cell types and incorporate a new set of relationships among them. Finally, we could strengthen the way we currently model the relationships between cell types (as a correlation matrix) by also adding higher-level categorizations – for example, through a feature that encodes which cell types belong to the B cell supertype, and similarly for T cell subsets.

The source of and modeling strategy for RNA-seq data deserves further consideration, as well. So far, we have trained *infino* on data from purified cell populations only. While this can provide a sense for the behavior of individual cell types, we incorporate no data suggesting how these cell types may mix. Incorporating some mixture training data could further aid prediction. However, little RNA-seq ground truth mixture data exists today, to our knowledge. The particular RNA-seq quantification strategy *infino* employs is another area of potential improvement. Since read counts are known to depend on transcript length, we could correct for this source of bias by adjusting for the length of each transcript [25]. There is also considerable debate in the literature as to which metric best quantifies transcript abundance. Other metrics, like FPKM, thus merit investigation.

Our approach would be even more useful with the ability to estimate the abundance of non-immune-cell content in the tumor microenvironment. Since samples are not uniform in their amounts of stromal tissue, immune cells, and tumor cells captured, controlling for this heterogeneity would enable better analysis. For example, by adjusting for the variation in immune cells between samples of two patients’ tumors, we could characterize the differential expression of the tumor cells. To estimate absolute abundances of immune cells in the microenvironment, rather than relative abundances as we have done so far, we propose incorporating a non-immune-cell component into our mixture model. That is, we could model a mixture as having immune, tumoral, and stromal components, or simply as having an immune component and a non-immune component. The non-immune-cell component could have a vague prior, or we can seed this “other” component with tumor cell lines. However, others have noted that these efforts may be complicated by the fact that tumor cells can sometimes mimic the expression patterns of immune cells, such as tumors with parainflammation exhibiting expression patterns characteristic of macrophages [15,26].

One technical challenge remains standing in the way of us applying *infino* to a large clinical dataset. Since the the joint distribution is modeled directly under the paradigm of Bayesian inference with a generative model, we estimate all model parameters simultaneously. As a result, the process of fitting *infino* and deconvolving ten unknown mixtures simultaneously routinely requires over two days of wall clock compute time for four simulation chains (parallelized). In this form, *infino* cannot practically score large collections of unknown mixtures.

We plan to investigate three modifications to our procedure for running *infino* intended to accelerate the process. First, we will evaluate the accuracy of variational inference methods, which bypass the lengthy simulation process and produce fast but noisy estimates. Variational inference could quickly provide a rough picture of the tumor microenvironment to a user, who could then choose to investigate further with a lengthier simulation by traditional methods. Additionally, variational inference is straightforward to integrate into our current infrastructure for running *infino*.

Second, the complexity of the training procedure depends on the number of genes we incorporate, since each transcript is represented by a set of parameters. While decreasing the number of genes used would lower the required time to deconvolve mixtures, the time savings would come at the expense of *infino’s* predictive accuracy. We will investigate how to choose a set of informative genes whose expression helps differentiate immune cell phenotypes, as well as a set of housekeeping genes with stable expression levels for a baseline.

Third, refitting parameters from scratch on every execution may be wasteful. While the Bayesian inference paradigm does not support a separation into “training” and “testing” phases, we can accomplish a similar separation of concerns by using stronger, more informative priors and supplying pre-fit hyperparameter values for these priors. This would effectively enable pre-computing model parameters related to our set of 63 training samples from purified cell populations. In particular, we recommend developing a solution to distribute *infino* by shipping informative priors. Rather than feeding in training data for every use, a user could instead supply a vector of parameter values estimated in an earlier offline run with the complete set of training data. This simple innovation could dramatically accelerate *infino* runs and enable evaluation of large clinical datasets -- bringing answers to consequential questions in cancer immunotherapy within reach.

## Methods

We propose a new method, *infino*, that enables improved diagnostics and clearer differentiation of similar cell types while capturing less noise and supporting RNA-seq data. Our approach is to deconvolve RNA-seq mixtures with a Bayesian generative model that encodes the process of mixing immune cell types. *Infino* estimates the probability distribution of each cell type’s expression profile, naturally incorporating variation and resolving a limitation of earlier approaches. Aggregating the posterior probability distributions at varying levels of the immune cell type hierarchy produces improved diagnostics for the evaluation of deconvolution results and performance. A critical innovation is the incorporation of relationships between cell types, which we demonstrated as storing valuable information capable of improving deconvolution accuracy (*Online Methods*).

## Online Methods

### Data

We obtained RNA-seq measurements from a publicly-available dataset of 63 purified immune cell populations [2]. To quantify the number of transcripts per gene, we created a RNA-seq processing pipeline using Google Container Engine and the Kubernetes technology [1,11] to replicate the processing described by [2]. We ran two Docker containers in series under massive parallelization through the batch job functionality of Google Container Engine’s hosted Kubernetes cluster service offering. The first container downloaded raw FASTQ RNA sequencing reads from a set of public dataset URLs. Then processing containers were run in parallel over the downloaded files, each one first unzipping the FASTQ reads, then performing trimming, which removes the bases with low quality reads – a commonly used but controversial technique [18]. Finally, the Kallisto tool was run over the preprocessed data to count the abundance of each gene transcript in the RNA-seq reads. The Kallisto tool is a popular choice because it avoids an expensive alignment step when quantifying transcript abundance [3]. In this manner, we downloaded the raw sequencing reads and executed standard quantification tools to clean the data and count the number of transcripts of each gene. The process ran for roughly an hour per sample.

### RNA sequencing transcript abundance transformation

Microarray modeling assumptions do not apply to the direct interpretation of RNA-seq data due to sampling bias and normalization requirements. We compared RNA-seq samples from the processed dataset [2] to similar microarray data to understand how to properly model RNA-seq mixtures. As suggested in [9], we log-transformed RNA-seq transcript counts.

In addition to different phenotypes being clearly distinguishable, even across technologies, the most highly expressed genes were observed to follow the same patterns. While microarrays have an independent probe for each transcript (in a limited set of transcripts), RNA sequencing pulls a finite number of reads from a pool of RNA. As a result, RNA-seq transcript counts are interdependent, since every read of one transcript leaves one fewer read for all other transcripts. Therefore, the most highly expressed transcripts may be expected to be in competition for the limited number of reads [14]. This suggests that the two most highly expressed genes, for example, will have a different relationship in RNA-seq data than in microarray data. Filtering to the highly expressed transcripts, we compared one transcript to the next most-expressed transcript. After applying the *voom* transformation, we found that microarray and RNA-seq data were comparable even in the relationships between pairs of very highly expressed genes. For instance, we observed a correlation of r=0.92 between the top 25 pairs of highly expressed genes.

### Generative modeling

In this study, we propose a Bayesian regression mixture deconvolution method that encodes the immune cell lineage relationships and produces rich confidence scores at all levels of the hierarchy of immune cell types. In this case, a generative model encodes the process by which mixtures are generated from the set of all immune cell types. Moreover, a Bayesian generative model can naturally incorporate the inter-cell-type relationships that we have called a “hierarchy”, and even learn these relationships directly from the data. We model expression and its variance for each cell type, not just a point estimate. Furthermore, we have the flexibility to condition on tissue of origin and similar predictors. The model will produce posterior probability distributions, which are much richer confidence scores than the point estimates and hypotheses tests used in earlier methods.

Importantly, generative models are distinct from discriminative models, which directly learn *P*(*y* | *x*), the probability of data *y* given predictors (independent variables) *x*. For example, in the classification context, discriminative models learn the decision boundaries between classes that are maximally “discriminative”, then use these boundaries to distinguish between the classes when labeling a test example. (For instance, support vector machines find an optimal hyperplane that represents the decision boundary separating classes.) Instead, generative models express a joint probability distribution over all observations and labels. This means they represent a full model incorporating all variables, including latent parameters. Generative models estimate *P*(*x, y*) first; rather than drawing decision boundaries between classes, generative models learn the distribution of each individual labeled class. Then to classify test examples, this joint distribution is transformed into *P*(*y* | *x*) by applying the definition of conditional probability: 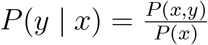. (The denominator *P*(*x*) is the empirical probability density of the data.) Again in the classification context, we can understand a generative model’s process for labeling a test example as computing which class is the most likely to have generated the data point, given that we know how data is generated because we have modeled each class’s distribution [13].

In the generative model paradigm which captures the process by which the data is generated, a researcher can estimate a latent (unmeasurable) variable, such as the mixture weights we seek that produce the observed mixture. We also state our prior beliefs, which can be vague for parameters such as the mixture components, or informative for each cell type’s expression distribution. Then we utilize a computational tool to repeatedly sample from the posterior joint probability distribution. At each step, we update our beliefs or uncertainty about the true mixture weights using Bayes’ rule, which follows directly from the definition of conditional probability and represents the fact that multiplying one’s prior belief with new evidence yield an updated belief distribution, called the posterior: *P*(hypothesis | data) ∝ *P*(data | hypothesis)*P*(hypothesis). This process is repeated until convergence of the posterior distribution [7].

### Stan probabilistic programming language

To express this Bayesian model and perform repeated sampling from the data generation process with updates to the posterior belief distribution at each step, we use Stan, a Turing-complete programming language in which random variables are first-class citizens. In particular, we use a python wrapper called *pystan*, one of several programming interfaces exposed by Stan [4]. A Stan program represents the conditional probability *P*(*θ* | *y, x*) of a generative model, where: *θ* are parameters, including the unknown latent mixture components; *y* are known data, such as the observed RNA-seq read counts; and *x* are predictors, including constants like the ground truth mixture weights for synthetic “training” mixtures. Then *P*(*θ, y, x*) is the joint probability over all data and parameters. *A priori* beliefs about the model, called “priors,” are encoded as *P* (*θ*).

After writing the generative model as a Stan program, it is compiled into a C++ executable that performs inference using a variant of Markov chain Monte Carlo sampling [8]. A set of query or input data is fed into the inference executable, which produces the requested amount of samples from the posterior joint probability distribution. Bayesian inference allows us to calculate the posterior *P*(*θ* | *y, x*), which represents the uncertainty in our beliefs of parameter values from our estimation with the available data. The joint distribution can be written as *P*(*θ, y, x*) = *P*(*y, x* | *θ*)*P*(*θ*), where *P*(*θ*) represents prior beliefs and *P*(*y, x* | *θ*) is the likelihood function *L*(*θ*) (a density of the observed data given the parameters) [10]. While exact inference is often impossible, the computation can be expressed as: 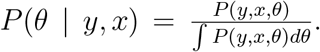. Finally, we can perform Bayesian predictive inference to evaluate the probability of a new observation *P*(*ỹ* | *y*) = *∫ P*(*ỹ* | *θ*)*P*(*θ* | *y*)*dθ*. In other words, given a new data point, we marginalize over the posterior to predict the new data point’s probability given the posterior distribution, which we estimated in the Bayesian inference step. This mechanism enables direct evaluation of how well the model fits our data [5,7,16].

The details of the sampling procedure are key to effective inference. Stan starts with initial guesses for parameter settings, then produces repeated simulated draws from the posterior distribution to correct the parameter posterior distributions until stationary distributions are reached. The initial samples are treated as warm-up iterations and excluded from the final results. To avoid reliance on those initial points, multiple chains of MCMC are used, and their convergence is evaluated [16].

### Bayesian model of RNA transcript mixture

We model gene expression mixtures as follows. Starting from a collection of cell types – each with their own expression distributions for all transcripts and with relationships to other cell types – we draw several cell types and mix them linearly with a specified weight for each cell type. This weighted average produces the transcript counts we observe in RNA-seq mixture data. Then we apply the inference machinery described above. By running multiple MCMC chains, we can detect whether there are multiple likely possible deconvolutions (in such a scenario, the separate MCMC sampling chains would not mix), affording a level of flexibility not available in existing approaches to immune infiltrate quantification.

Our model incorporates the following features:

- Estimated counts for each transcript in every sample.
- For each cell type, a per-gene offset from that gene’s mean expression level across all cell types and samples.
- A correlation matrix between the above offsets for each cell type to incorporate cell type relationships.
- A scale for each cell type that is multiplied on diagonal with the correlation matrix to form the covariance matrix between cell types. (This forms a hierarchical model of relationships among cell types.)
- Cell-type specific predictors, including surface markers.
- The weight of each cell-type specific predictor across cell types.
- For each transcript, an overdispersion parameter across all samples to account for RNA-seq read count heteroscedasticity, estimating variance in transcript-level expression among samples.
- An adjustment for the expression level of “housekeeping genes.”

Within the probabilistic programming paradigm, our “query data” includes:

- Training data: single-origin purified cell population samples of known composition (e.g. entirely naive B cells).
- Testing data: mixtures of unknown composition.

The model infers sample compositions whose fractional components sum to one. To do so, the model first estimates the relative expression offsets of each transcript in each cell type. This is a standard multivariate regression problem, therefore we apply the multivariate normal distribution, which is the Gaussian distribution extended to a high-dimensional vector.

The model then estimates the pairwise correlation matrix between the cell types. This can be viewed more precisely as a distance computation, encoding how different the expressions of each cell type are. From this perspective, the correlation matrix is a broad way to represent a hierarchy of cell types, because it places similar cell types closer together without enforcing the rigidity of a hierarchy expressed in tree form. That is, a tree hierarchy assumes that certain relationships cannot exist across lineages, for instance. Instead, the correlation matrix representation of cell-type similarity is more general and enables more relationships to be learned. We will hereafter refer to our modeling innovation as using a correlation matrix rather than a hierarchy. In fact, “hierarchical Bayesian modeling” generally refers to multilevel modeling with a hierarchy of features (parameters), rather than a hierarchy of labels (cell types). We note that our model is also multilevel, since there is a hierarchy among the parameters.

Next, the model estimates the observed expression in the training and testing samples. The mixture is represented as a constant base expression level for each gene, to which cell-type specific offsets mixed with fractional weights are added – a master distribution with deviations. Those weights are the desired deconvolution fractions. Transcript counts are modeled as a gamma-Poisson distribution mixture, which is the negative binomial counts distribution. This is essentially a Poisson count distribution, with a gamma distribution underlying it to account for variability and overdispersion. As noted by [9], the heteroscedasticity of transformed RNA sequencing read counts must be considered; hence, we allow for overdispersion and per-transcript variability beyond a standard Poisson counts model. While transcript counts are fed into the model in their raw form (because log-transformed counts do not follow a normal distribution), we apply a log link function in the negative binomial distribution to effectively model log-transformed counts, per [9].

We specify our hierarchical generalized linear model [7] as follows:

We relate a linear predictor, *Xβ*, to our outcome variable matrix of gene counts ***Y***, a (*S* × *G*) matrix where entry *Y_s,g_* corresponds to the mean number (tpm) of transcripts of *g* in sample *s. X* is a (*S* × *C*) matrix corresponding to the abundance of cell type per sample, and *β* is a (*C* × *G*) matrix representing the gene transcript counts per cell type.

We relate the expectation of our outcome variable *y* with our linear predictor like so:

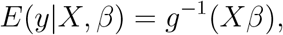

where the “link function” *g* is a negative binomial parameterized by *μ*, the log of the mean expression per gene, and *ϕ*, the dispersion parameter per gene. Then:

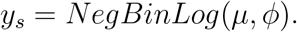

For each sample, *μ* (a vector of *G* elements) is further decomposed into a transcript-level log-mean 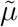 (a vector of *G* elements), cell-type-specific transcript abundances *β*, and the sample composition 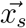, where *x_s_i__* ∈ [0, 1]∀*i* ∈ [1, *C*] and 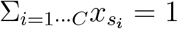:

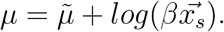

We place a hyperprior on our coefficient *β* with a multinormal on each gene *g*’s vector of expression per cell type:

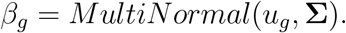

For a gene *g*, the vector of cell-type-specific means *u_g_* has contributions: from the mean expression level of the gene per cell type, *p*; the *C* × *M* matrix of cell-type features, *F*; the coefficients per feature *b* and each feature’s influence on a gene *κ*. That is:

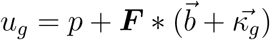

Finally, the covariance matrix Σ is decomposed into the diagonal matrix (“scaling factor”) *τ* and correlation matrix Ω:

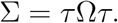

It is the correlation matrix Ω that our model computes as encoding the hierarchy of immune cell types.

### Validation of model convergence

When measuring the model’s predictive accuracy with synthetic mixtures, we evaluated the model’s convergence. We generated synthetic test mixture of known composition from combinations of the 63 purified cell populations. Then we fit *infino* with only the expression mixture and not the mixture fractions. That is, we supplied only the simulated expression values, not the fractional mixture components, to the model, to assess the model’s estimated mixture components.

We fit the model with the NUTS sampler through *pystan*. The model fit lasted 51 hours and 20 minutes to produce four MCMC chains of 2000 iterations each (in parallel). The first 1000 samples of each chain were considered to be warm-up samples and discarded.

First, we checked whether convergence was reached in the sampling. We plotted the Monte Carlo standard error, which measures the consequences of a limited number of sampling draws (Figure 1). We observe that the Monte Carlo standard error was under 2%, an order of magnitude lower than the highest posterior standard deviation of a parameter estimate (Figure 2). This suggests that our sampling strategy was effective [5,7].

**Figure 1:**
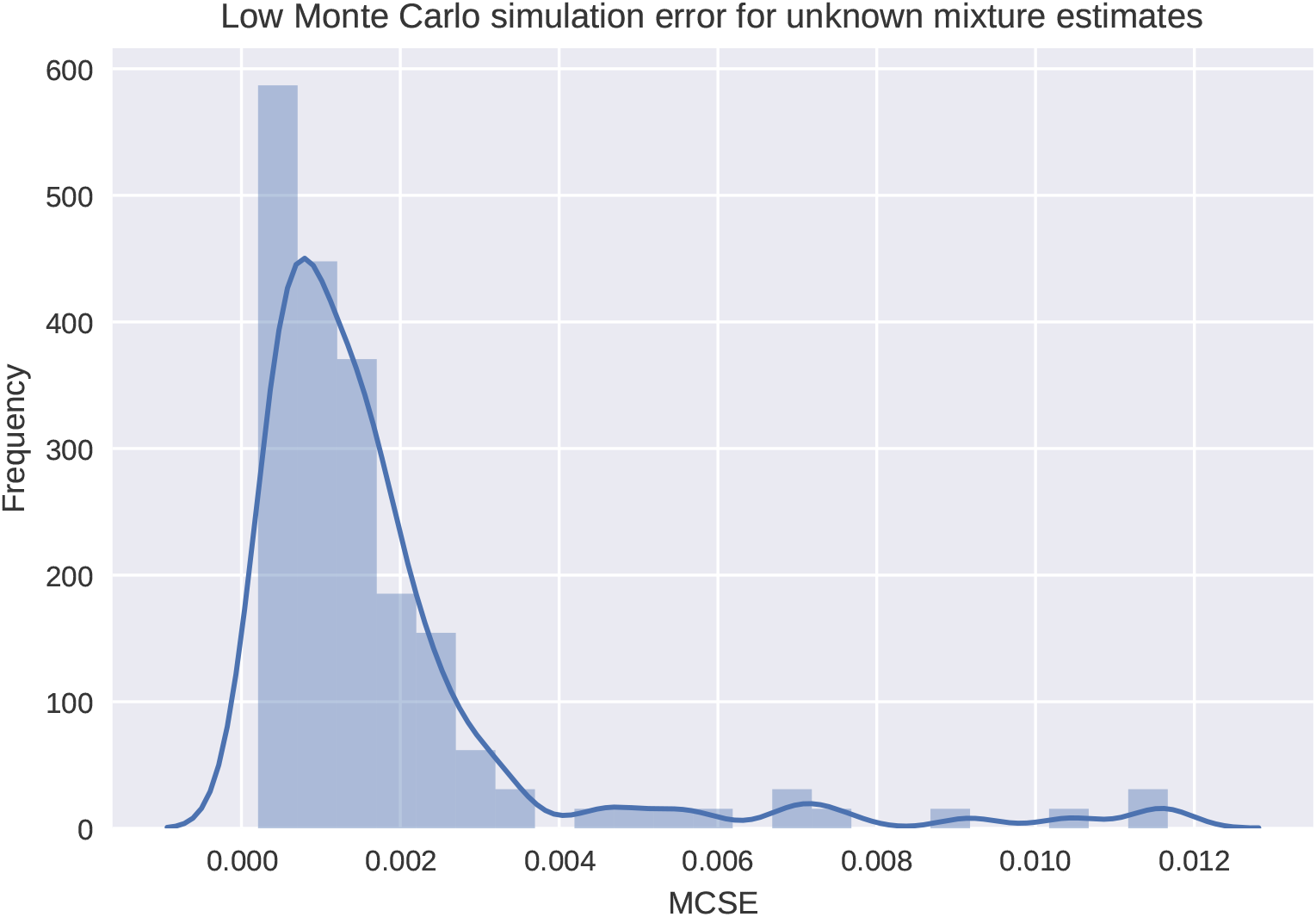
A histogram of the Monte Carlo standard errors of all unknown mixture component parameter estimates.

Second, we examined the level of autocorrelation in our sampling. By definition, MCMC is serially correlated: each parameter configuration is a random deviation away from the previous parameter configuration. Ideally, the level of correlation between successive samples is low. We plotted a histogram of the distribution of effective sample sizes across all parameter estimates for the unknown mixture components (Figure 3). In this sampling run, the effective sample size was quite variable, but there were always at least a few dozen usable samples per parameter, suggesting that we sampled enough to trust the model’s estimates.

**Figure 2:**
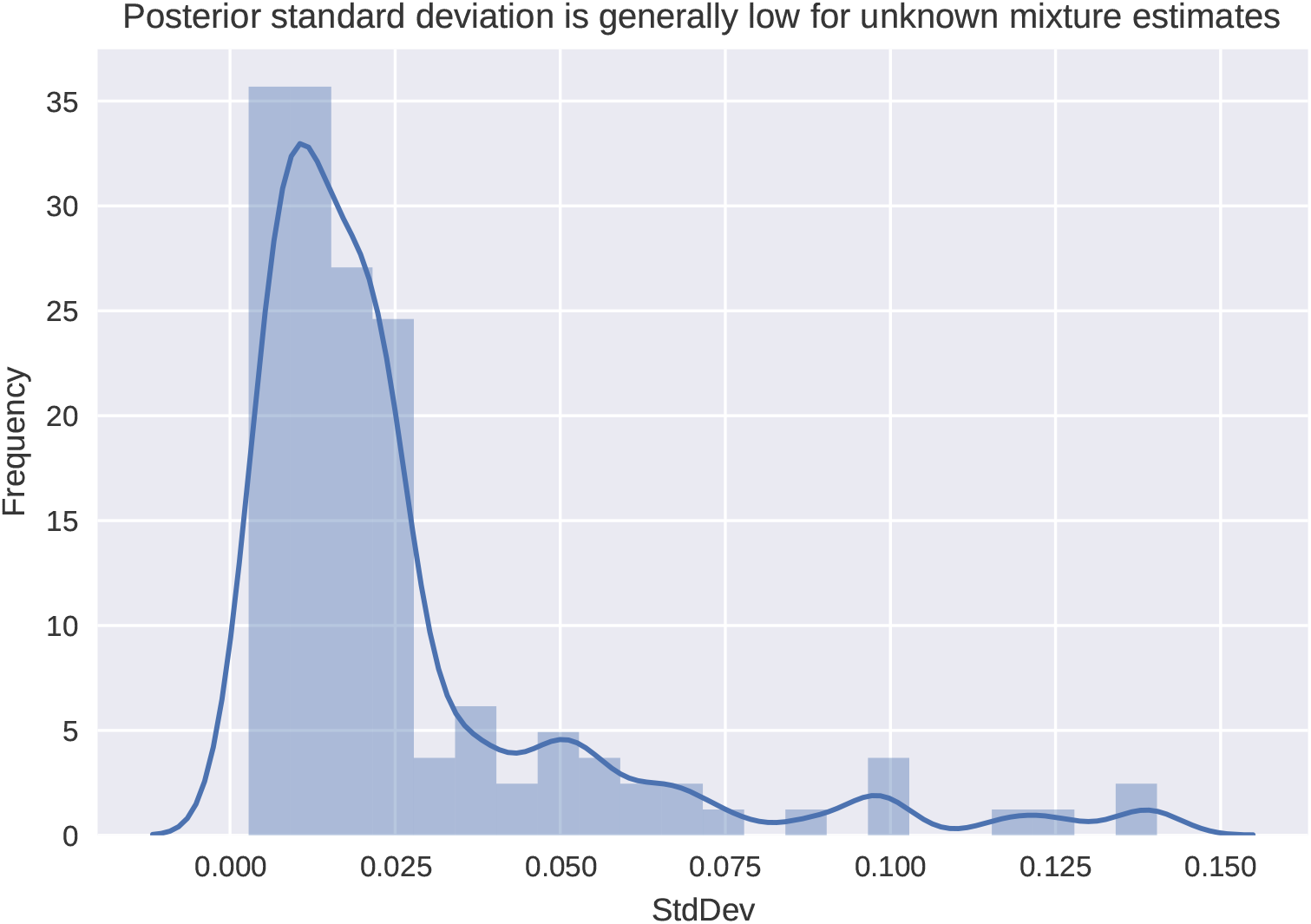
A histogram of the posterior standard deviations of all unknown mixture component parameter estimates.

**Figure 3:**
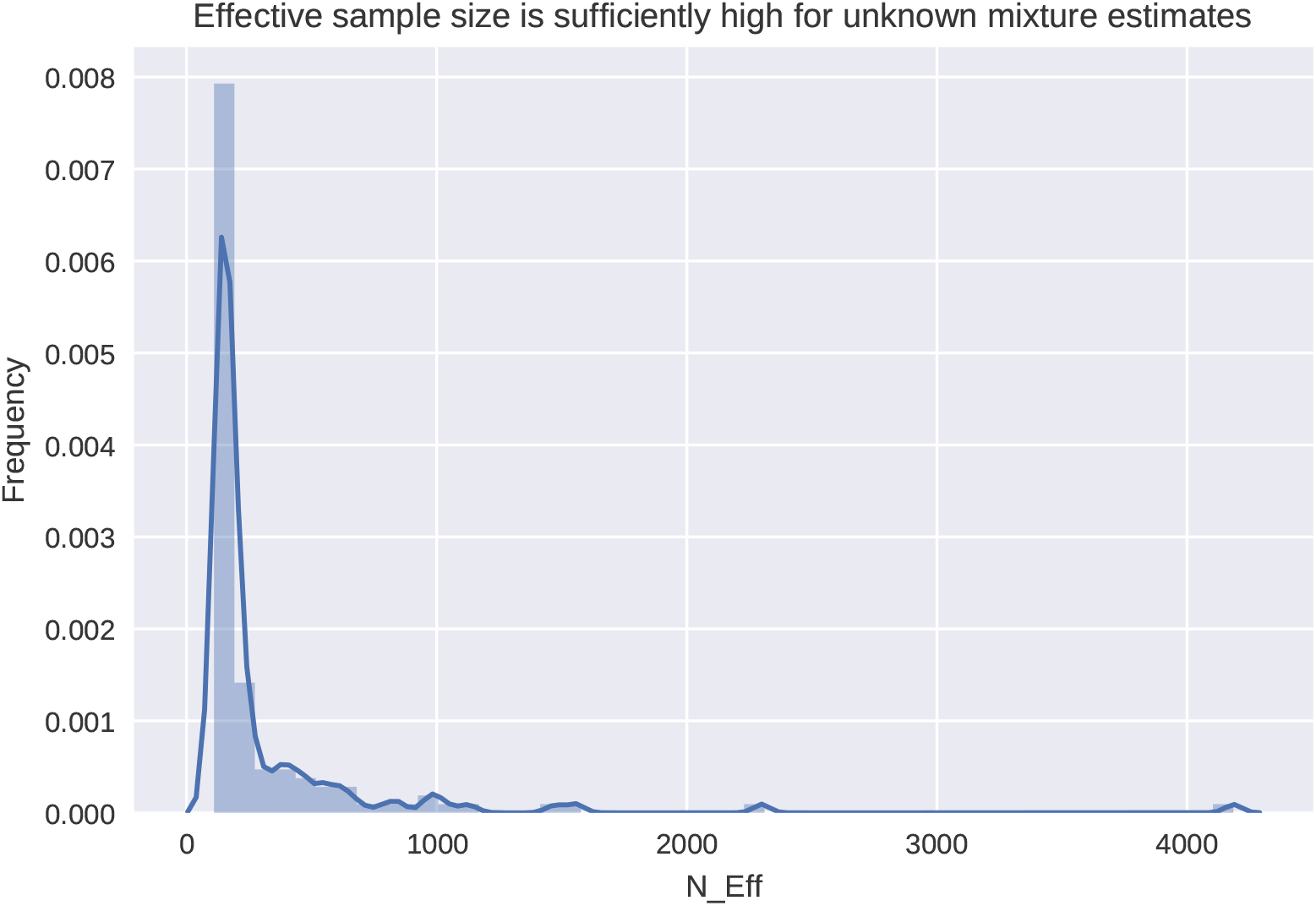
A histogram of the effective sample sizes of all unknown mixture component parameter estimates.

We also examined the parameter traceplots, which show how the four chains sampled a parameter in all of their iterations. Figure 4 displays the simulation trace of the posterior estimate of one unknown mixture component. This traceplot reveals low autocorrelation, since the parameter estimate jumps widely rather than shifting slowly [5,7].

**Figure 4:**
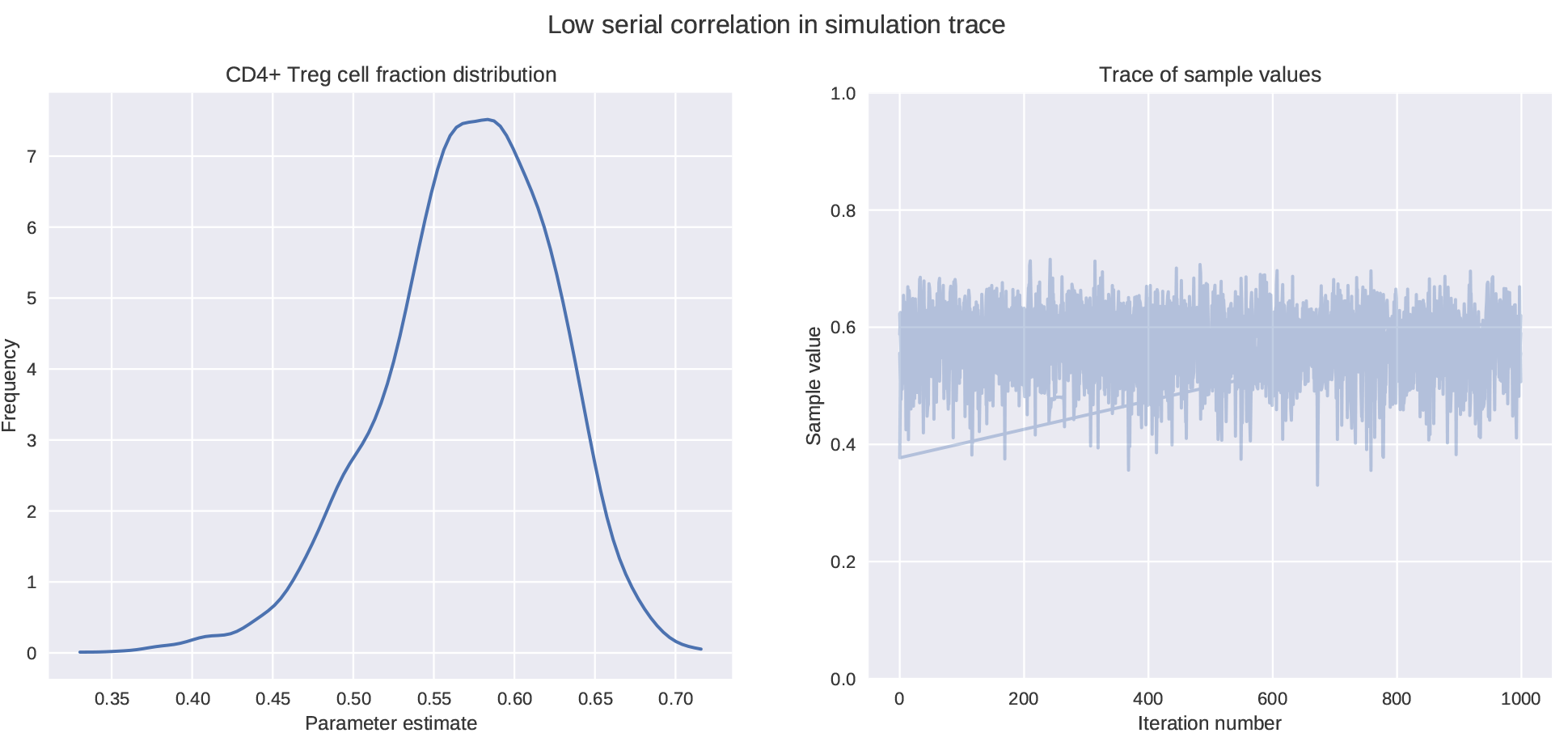
The trace of the CD4+ Treg cell estimated fractional mixture weight component in a 25%-75% mix of naive B cells and CD4+ Treg cells.

The traceplots also demonstrate that the four chains mixed quite well, since they appear indistinguishable. This suggests that the unknown mixtures did not have multiple valid deconvolutions discovered by separate MCMC chains. Rather, only one deconvolution for these simulated mixtures appears to be valid.

We can also look to 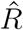, the Gelman-Rubin potential scale reduction factor, which is a metric of how well the chains converge [6]. Since the 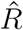 values for the unknown mixture fraction parameters were close to one (Figure 5), we conclude that variation in the chains’ estimates of these parameters would not be reduced significantly by longer chains [5,7]. Also, we note that the 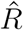 for the estimated log posterior likelihood was low (1.0049), suggesting overall model convergence was achieved.

**Figure 5:**
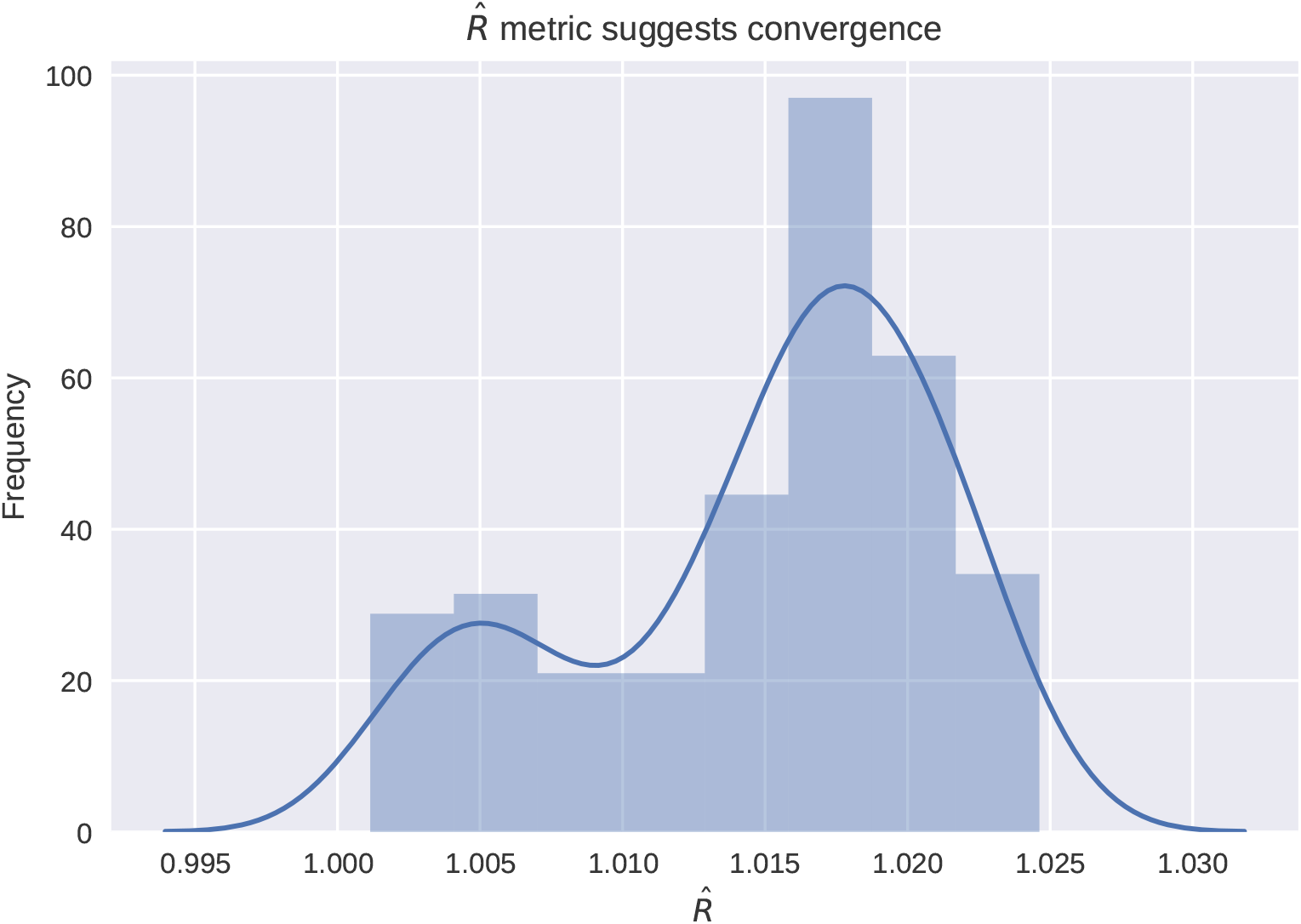
A histogram of the Gelman-Rubin potential scale reduction factors of all unknown mixture component parameter estimates.

These tests suggest that the estimated posterior distributions converged in our simulation run and their variance would not be significantly decreased by sampling further. Therefore, we are confident that these results represent the model’s true ability to deconvolve, and now will analyze how the model learns expression data and cell type relationships.

### Hierarchical clustering of estimated correlations between cell types

First, we extracted the posterior distribution samples for the correlation matrix parameter in our model. We then converted every correlation *r* into a distance metric *d* as follows: 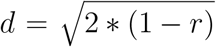. We employed hierarchical clustering to visualize the estimated hierarchy in the form of a dendrogram. Specifically, we progressively grouped cell types into clusters by their pairwise distances using Ward’s method, which minimizes variance within created clusters [17]. The cophenetic correlation, a measure of how accurately a hierarchical clustering dendrogram represents the true pairwise distances, was 0.910 for our clustering, where 1 is best [15]. Therefore, we trust that the clustering preserves the true relationships in the estimated correlation matrix.

### Source code

Instructions for running *infino* are available at https://github.com/hammerlab/infino.

### Comparison to Cibersort

All comparisons were performed using `Cibersort v1.03` [12] and default settings as described in the Cibersort documentation.

